# Paraoxonase and acylated homoserine lactones in urine from patients with urinary tract infections-- relationship to microbial diversity by 16S rRNA gene sequencing

**DOI:** 10.1101/2021.06.10.447923

**Authors:** John Lafleur, Jacquelyn S. Meisel, Seth Commichaux, Richard L. Amdur, Mihai Pop, Mark W. Silby

## Abstract

Paraoxonase (PON) comprises a trio of mammalian enzymes that have been reported to have a number of roles including the inhibition of bacterial virulence and biofilm formation by microorganisms that quorum sense with acylated homoserine lactones (AHLs). PON have previously been reported to inhibit *P. aeruginosa* biofilm formation in mammalian airways and skin. An innate immune role for PON in urinary tract infection has not previously been reported. We performed western blots for PON1 in urine from patients with urinary tract infection (UTI), and also tested UTI urine for the presence of AHLs using a cellular reporter system. Urine sample microbiota was assessed through sequencing of the 16S rRNA marker gene. We report here that PON1 was not found in the urine of control subjects, however, in patients with UTI, PON1 was associated with the presence of *E. coli* in urine. AHLs, but not PON, were found in the bulk urine of those with *P. aeruginosa* UTI. Microbial consortia of PON positive UTI urine was found to be distinct from PON negative UTI urine; differentially over-represented bacteria in PON positive samples included a number of environmental opportunists. We hypothesize that PON may inhibit the quorum sensing activity of AHLs in UTI, as has previously described in skin and airways.

## Introduction

The paraoxonase family (PONs) of mammalian lactonases are an evolutionarily conserved (1–3) innate immune mechanism that limit bacterial virulence and biofilm formation by degrading quorum sensing (QS) acylated homoserine lactones (AHLs) produced by some bacteria (4–13). These AHL-producing bacteria include *P. aeruginosa,* as well as other environmental opportunists with large genomes and flexible lifestyles that are frequently found to be occult members of infecting biofilms (14–18). UTIs caused by such organisms have been reported to be more common in patients with urinary catheters, diabetes mellitus, and previous hospitalizations (19). In multiple studies PONs have been shown to be protective against infection by *P. aeruginosa* biofilms in mammalian airways and skin cells (9, 20). Another body surface that is subject to environmental exposure is the urinary system, and PON1 has previously been reported in minute concentrations in specialized vesicles in urine from healthy subjects (21).

AHLs are known to be potent activators of quorum sensing that favors biofilm formation and virulence gene expression in certain gram negative bacteria (22). In parallel, AHLs can directly induce tissue inflammation and derangement of host immunity (23–27). AHLs have not previously been reported in urine from human UTI although their presence would have significant implications for the diagnosis and treatment of UTI.

Uncomplicated UTI in an immunocompetent host is characterized by the predominance of a single bacterial species (28), *E. coli* in 80% of cases. One possibility is that the predominance of *E. coli* in uncomplicated UTI is due to innate immune activity toward opportunists/difficult to eradicate environmental strains. In particular, we speculated that environmental opportunists, such as *P. aeruginosa,* are inhibited from quorum sensing with AHLs due to PON activity in the urine. PON has been shown to limit *P. aeruginosa* virulence in Drosophila—a model eukaryotic organism that does not naturally produce PON (7). PON-deficient mice display increased vulnerability to infection with *P. aeruginosa* (29). Whether and how PON is induced in mammalian infection is not at present known in detail, however protection against *P. aeruginosa-*mediated virulence has been shown to involve induction of PON2 through peroxisome proliferator-activated receptor-g (30, 31) which is found in mammalian urothelial cells (32), though the AHL QS molecule (3OC12-HSL) produced by *P. aeruginosa* has been shown to have inhibitory effect on this receptor (33). Another possibility is that PON are induced through Toll-like receptors (TLRs) which are cellular sensors for a variety of bacterial factors. Urinary TLR signaling has been found to be sensitive to uropathogens (34, 35), resulting in activation of NF-kB and the expression of the pro-inflammatory genes IL-6 and IL-8 with consequent ingress of neutrophils to the bladder mucosa (36). There is at present, however, no report in the literature of TLR mobilization of PON. Host disruption of AHL QS through induction of PON has been proposed as protective against inflammatory bowel disease (37). While urea-mediated inhibition of QS mechanisms in chronic *P. aeruginosa* infection have been shown to limit biofilm formation and other virulence factors without inhibiting the production of AHLs in a murine CAUTI model (38), in acute UTI with *P. aeruginosa*, AHL QS has been shown to promote virulence (39). In addition, it has been shown that PON mediates changes in microbiota in the Drosophila gut (40), and in a flow cell model of a polymicrobial consortia (41). In summary PON are an effector of innate immunity (29, 42) that inhibit bacterial QS-mediated virulence through degrading the AHL QS signal. In addition, PON have been shown to alter the composition of host microbiota. To explore possible implications of this in the urinary system, we set out to measure AHLs and PON in urine from patients with UTI presenting to an emergency department of an urban hospital. We also assessed the microbiota of study samples through 16S rRNA gene sequencing. We hypothesized that 1) UTI patients will have PON present, while non-UTI patients will not; 2) among patients presenting with urinary symptoms (dysuria and frequency) and urinalysis showing elevated urine leukocytes, the presence of PON will be associated with growth of a urinary pathogen in culture; 3) UTI urine with PON will be associated with urinary microbiota distinct from that of UTI urine without PON.

## Materials and methods

### Human subject enrollment

Study protocol was reviewed/approved by the Miriam Hospital IRB (#496193-16). Written informed consent was obtained. The study was conducted at the Anderson emergency department of Rhode Island Hospital. Inclusion criteria for entry into the study: Greater than 18 years of age and able to give informed consent for study participation; 10 or more white cells in urine analysis with symptoms of urinary tract infection; urine culture sent to the hospital microbiology department (prior to administration of antibiotics). Control subjects were emergency department patients with minor complaints unrelated to urinary system and without significant metabolic derangement such as fever, hyperglycemia, renal disease (acute or chronic), or significant hypertension. Once enrolled study subjects were asked to provide 50-100 ml of clean-catch urine in a sterile cup. This was immediately frozen at −80 for further study. Culture results and clinical data were obtained through the electronic medical record at Rhode Island Hospital.

### PON and AHL assays

#### Growth media

Plates and broth were lysogeny broth (LB).

#### Strains

The long chain HSL reporter strain *E. coli* JM109 (pSB1142) (carries *P. aeruginosa las*R and the *lasI* promoter fused to *luxCDABE*) (43), and *P. aeruginosa* PAO1 carrying P_*lasB*_-*luxCDABE* (44) were grown in LB broth with shaking at 37 deg. C.

#### Reagents

3-oxo-C12-HSL stock solution 20 mg/ml (Cayman Chemicals) was diluted to 4 micrograms/milliliter in water. Dilutions were arrived at empirically by testing against luminescence in the long chain HSL reporter strain *E. coli* JM109 (pSB1142).

#### Western blotting

Urine samples from enrolled research subjects with UTI were stored at −80 deg. C., and thawed for use. 25 microliter samples of unprocessed urine were assayed for PON1 using the Bio-Rad iBlot system as previously described (45).

#### Antibodies

Primary antibody: polyclonal human PON1 from rabbit (HPA001610, Atlas Antibodies). Secondary antibody: goat anti-rabbit labeled with peroxidase (Invitrogen)

#### AHL assay

Construction of standard curve for 3-oxo-C12-HSL was determined by varying concentrations of the stock solution diluted in water and incubated with *E coli* JM109 (pSB1142), using a microtiter plate reading format(46) as previously described (47). 3-oxo-C12-HSL concentrations in samples were determined by adding 50 microliters of study urine samples to 5 microliters of overnight sensor strain *E. coli* JM109 (pSB1142) and then measured in a Varioskan Flash plate reader.

#### Urine Microbiome

Urine samples, (previously stored at −80 deg C.) in 50 ml quantities, were centrifuged at 2000 rpm for 20 minutes. Supernatant was poured off, and pellet was resuspended in 1 ml of sterile water. DNA was extracted using the FastDNA SPIN Kit (MP Biomedicals) (48)

### Preparation of 16S rDNA amplicon inserts for Next-Generation library construction and NGS sequence analysis using sequential PCR amplification steps

Sample preparation and sequencing was performed at the at the UMASS core facility in Shrewsbury, MA. 16S PCR was initially performed to add indexes to individual templates. 10 microliters of DNA template (10 nanograms) were amplified with primers for 16S V1V2 hypervariable region (figure 1—all primers were added in 1 microliter volumes from 10 micromolar stock solutions) (49) with Platinum PCR Super Mix (1306, LifeTech). 45 microliter reaction mixtures were placed in the wells of MicroAmp Fast Optimal 96-well Reaction Plates (0.1microliter) and run on a 7500 ABI Fast Real-Time PCR System with the cycling parameters: (95◻ 2 min) + 22 x (95◻ 45 sec, 50◻ 45 sec, 72◻ 1 min) + (72◻ 7min) + 4◻ O/N. Reaction clean-up was performed with Qiagen 96-well PCR cleanup plates; and PCR Product quantitated and profiled using an Advanced Analytics DNA Fragment Analyzer, Qubit, and NanoVue. A second 16S PCR was performed to add NGS adapter to barcoded templates with the same protocol (Table 1). Samples were sequenced on an Illumina MiSeq using 300 bp PE chemistry. Reads were processed and amplicon sequence variants (ASVs) were generated using DADA2 in R. Reads were quality trimmed and filtered using the command fastqPairedFilter using parameters trimLeft=c(10, 20), truncLen=c(240, 200), maxEE=2, rm.phix=TRUE, rm.lowcomplex=5, kmerSize=2. DADA2 was used to learn error rates, perform sample inference, dereplicate and merge paired end reads, and construct a sequence table. Taxonomy was assigned using the SILVA 132 ribosomal RNA (rRNA) database.

**Table 1.**
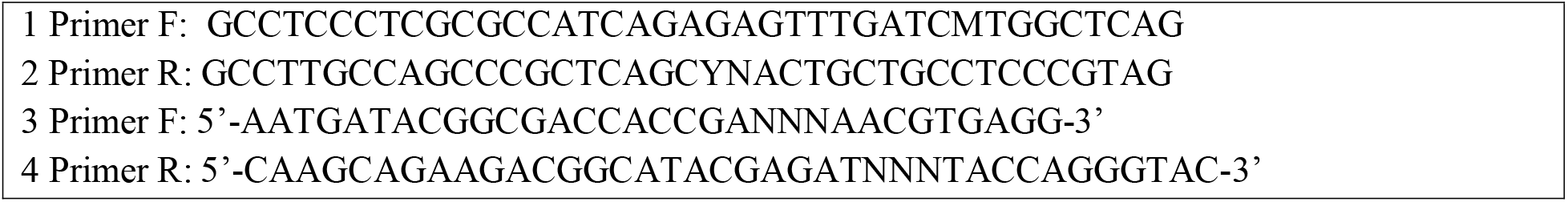
Primers 1&2 were used in the first PCR to add barcode; 3&4 in the second PCR reaction to add NGS adapters

**Figure 1.**
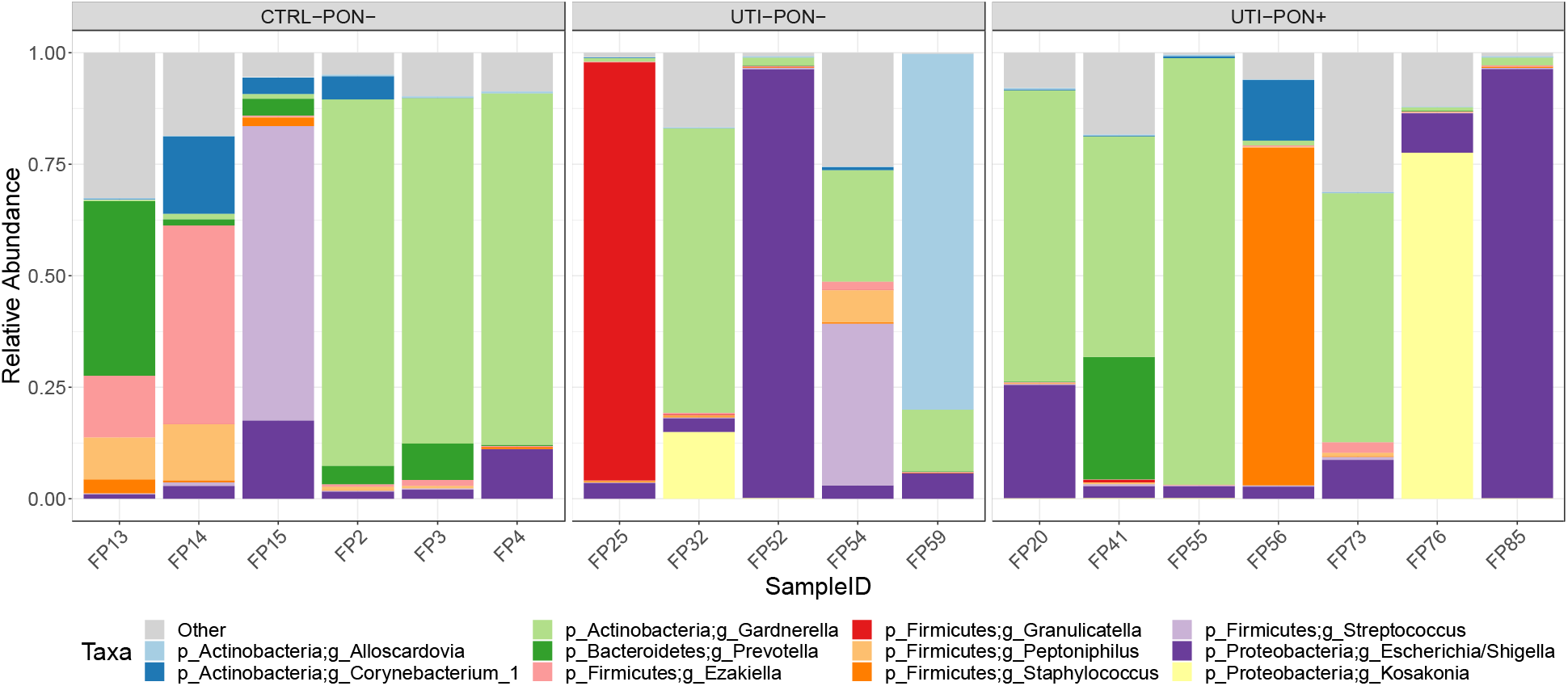
Bars show color-coded proportions of the top 12 taxa in urine microbiome samples, grouped according to UTI and PON status.

### Measures

Culture results were recoded into a binary variable (positive/negative). As a sensitivity analysis, we also coded those patients who were positive but with <50k cfu as negative. Positively skewed continuous variables and those with outliers were recoded into ordinal variables.

### Data analysis

Associations between diagnosis and categorical variables were analyzed using chi-square or Fishers Exact Test. Comparison of continuous variables across groups was done using 2-tailed independent groups t-tests or the Kruskal-Wallis test for skewed variables. In order to test the independent association of PON1 with being culture-positive in UTI patients, we used multivariate logistic regression, adjusting for variables that might be confounds. These were defined as having an association with PON1 antigen with p<.10. We also used a multivariable logistic regression model to develop an optimal prediction model for being culture-positive in UTI patients. This was based on the patient variables that were associated with being culture-positive with p<.10, dropping any for which an odds ratio could not be calculated due to low sample size. SAS version 9.4 (Cary, NC) was used for data analysis, with p<.05 considered significant.

### Microbiome analysis

A total of 24 samples had sufficient PCR amplification to pass quality control. These were processed for further analysis with bioinformatics tools as previously described. [ref.] Of these, 2 were technical controls, and 4 samples were found to have insufficient sequencing depth (< 1,000 reads per sample) and were not included in the analysis. Of the remaining 18 samples, 6 were controls, 5 were UTI/PON−, and 7 UTI/PON+. These 18 samples had a median sequencing depth of 132,986 reads. A total of 208 OTUs observed more than 3 times in at least 20% of the samples were retained for analysis in R using the packages phyloseq (50), breakaway (51), DivNet (52), and corncob (53).

## Results

### PON and clinical parameters

Mean age of enrolled subjects was 60 ± 22, 13 (19%) were black and 44 (63%) were white, 48 (69%) were female, and 11 (16%) had urinary catheters. Culture was positive with one or more uropathogens in 39/61 cases (64%), while western blot for PON1 antigen was positive in 22 cases (36%). There were 61 UTI patients and 9 controls in the sample.

Controls and UTI patients differed significantly in age, serum creatinine, highest temperature, and lowest diastolic blood pressure (Table 2.). Patients with UTI had higher average creatinine, likely due to age-related decline in kidney function; higher temperature in UTI subjects is likely due to some subjects being systemically ill.

**Table 2.** Patient variables by diagnosis (UTI vs control)

| Patient variable | Control (n=9) | UTI (n=61) | p |
| --- | --- | --- | --- |
| Age | 42 ± 16 | 63 ± 22 | .009 |
| Race |  |  | .45 |
| Black | 3 (33%) | 10 (16%) |  |
| White | 5 (56%) | 39 (64%) |  |
| Other | 1 (11%) | 12 (20%) |  |
| Gender female | 4 (44%) | 44 (72%) | .13 |
| Catheterized | 0 | 11 (18%) | .34 |
| WBC | 9.3 ± 4.4 | 11.4 ± 4.4 | .25 |
| Hemoglobin | 13.7 ± 0.9 | 12.5 ± 1.9 | .07<sup>A</sup> |
| Serum creatinine | 0.8 ± 0.1 | 1.2 ± 0.9 | .042<sup>A</sup> |
| Highest HR | 85 ± 14 | 93 ± 21 | .29 |
| Highest temp | 97.7 ± 0.6 | 99.0 ± 1.5 | .0001 |
| Lowest systolic bp | 123 ± 16 | 118 ± 20 | .40 |
| Lowest diastolic bp | 78 ± 11 | 68 ± 13 | .044 |

<sup>A</sup> using Kruskal-Wallis test.

PON1 was significantly associated with UTI diagnosis. Of the 61 UTI patients, 22 (36%) were PON1 positive, while none of the controls were PON1 positive (Fisher Exact test p=.049). PON1 was not significantly associated with any demographic or laboratory values (Table 3), but was significantly associated with higher heart rate (HR) (Table 3).

**Table 3.** Associations between patient variables and PON1 in patients with UTI

| Patient variable | PON1 Neg (n=39) | PON1 Pos (n=22) | p |
| --- | --- | --- | --- |
| Age | 62 ± 22 | 63 ± 21 | .82 |
| Race |  |  | .60 |
| Black | 5 (13%) | 5 (23%) |  |
| White | 26 (67%) | 13 (59%) |  |
| Other | 8 (21%) | 4 (18%) |  |
| Gender female | 29 (74%) | 15 (68%) | .61 |
| Catheterized | 5 (13%) | 6 (27%) | .18 |
| WBC | 11.2 ± 4.4 | 11.7 ± 4.6 | .68 |
| Hemoglobin | 12.5 ± 2.0 | 12.4 ± 1.7 | .75 |
| Serum creatinine | 1.2 ± 1.0 | 1.2 ± 0.5 | .48 |
| Highest HR | 88 ± 16 | 101 ± 25 | .03<sup>A</sup> |
| Highest temp | 99.0 ± 1.6 | 98.9 ± 1.3 | .66 |
| Lowest systolic bp | 120 ± 20 | 114 ± 20 | .27 |
| Lowest diastolic bp | 68 ± 12 | 70 ± 13 | .45 |

<sup>A</sup> using Kruskal-Wallis test.

PON1 was significantly associated with positive culture in UTI patients: PON was positive in 4/22 with negative culture (18%) versus 18/39 with positive culture (46%; p=.03; Fishers Exact test; Supplemental Table 1). We did a sensitivity analysis in patients who had culture < 50,000 cfu (culture negative), and found that the association was still significant (culture negative had 23% PON positive, culture positive had 48% PON positive, p=.04). Thus, in UTI patients, presence of PON in urine was associated with urine culture growing out a urinary pathogen, in contrast to urogenital flora, or no growth.

In addition to PON1, other patient variables that were associated with being culture-positive, in UTI patients, included WBC (higher with culture positive), and being catheterized (more frequent for culture positive). Highest HR was marginally associated with culture-positive (Table Supplemental Table 1.). To demonstrate the potential clinical utility of PON measurement in UTI we created an optimal prediction model for being culture-positive, which included PON1, highest HR, and WBC (being catheterized was dropped because OR could not be calculated for this variable due to small sample size), which had an area under the ROC curve of 0.72 for predicting culture-positive.

Using the equation: risk = −2.43 + 1.007*PON1 + .014*highestHR + .138*WBC, and then probability = exp(risk) / (1 + exp(risk)). Splitting the probabilities into tertiles, we found that the observed incidence of being culture positive in tertiles 1 through 3, respectively, were 39%, 75%, and 83% (p=.01).

Half of all positive urine cultures grew out *E. coli* alone (19/38). PON1 was positive in 10 of these (53%). When compared to cultures that grew out multiple organisms (including those with ‘urogenital flora’), PON was significantly associated with cultures that grew out *E. coli* alone, P=0.05 (Supplemental Table 2.). Seven gram-negative environmental opportunists were cultured from PON negative urines. Four were *P. aeruginosa*; the others were*: Serratia marscesens, Citrobacter freundii,* and *Klebsiella pneumoniae*. These organisms, like *P. aeruginosa*, are multi-drug resistant environmental opportunists. Additionally these bacteria have all been reported to produce or QS with AHLs.(54–56) Among PON+ urines no such taxa grew out in culture, though, gram negative environmental opportunists were over-represented as members of the urinary microbiome of PON+ urines (Figure 3).

### Measurement of AHLs in urine samples

Using an *E. coli* luminescent reporter construct (*E. coli* JM109 (pSB1142)), the presence and abundance of long chain AHLs was also assayed in urine samples. Long chain AHLs were only detected in three out of four urine samples that were culture positive for *P. aeruginosa* and PON1 negative. AHL concentration in one sample (patient #9) was about 1.5 micromolar. The other three samples in which *P. aeruginosa* grew out of culture had considerably lower concentrations (see Supplemental Table 3.).

### Urine microbiome

Similar to other human-associated microbiome studies (57), the taxonomic composition of the samples varied widely across individuals (Figure 1). Shannon diversity, which accounts for the richness and evenness of taxa within samples, was different between controls and UTI samples, but not between PON+ and PON− UTI samples (Figure 2A, p-value < 0.05). Similarly, Bray Curtis dissimilarity revealed greater differences between samples based on UTI status, than between PON+ and PON− UTI samples (Figure 2B). A total of 22 taxa were significantly differentially abundant between PON+ and PON− samples when controlling for differences based on UTI status (Figure 3). Those associated with PON− were typical host commensals such as *Corynebacterium* and *Peptoniphilus spp*.; all were gram positive cocci. Among the PON+ group, there was a more diverse group represented including commensals typically associated with mammalian hosts, and environmental bacteria found in a variety of ecosystems such as *Caulobacter* and *Aerococcus* which have both been found to cause human infection (58,59).

**Figure 2.**
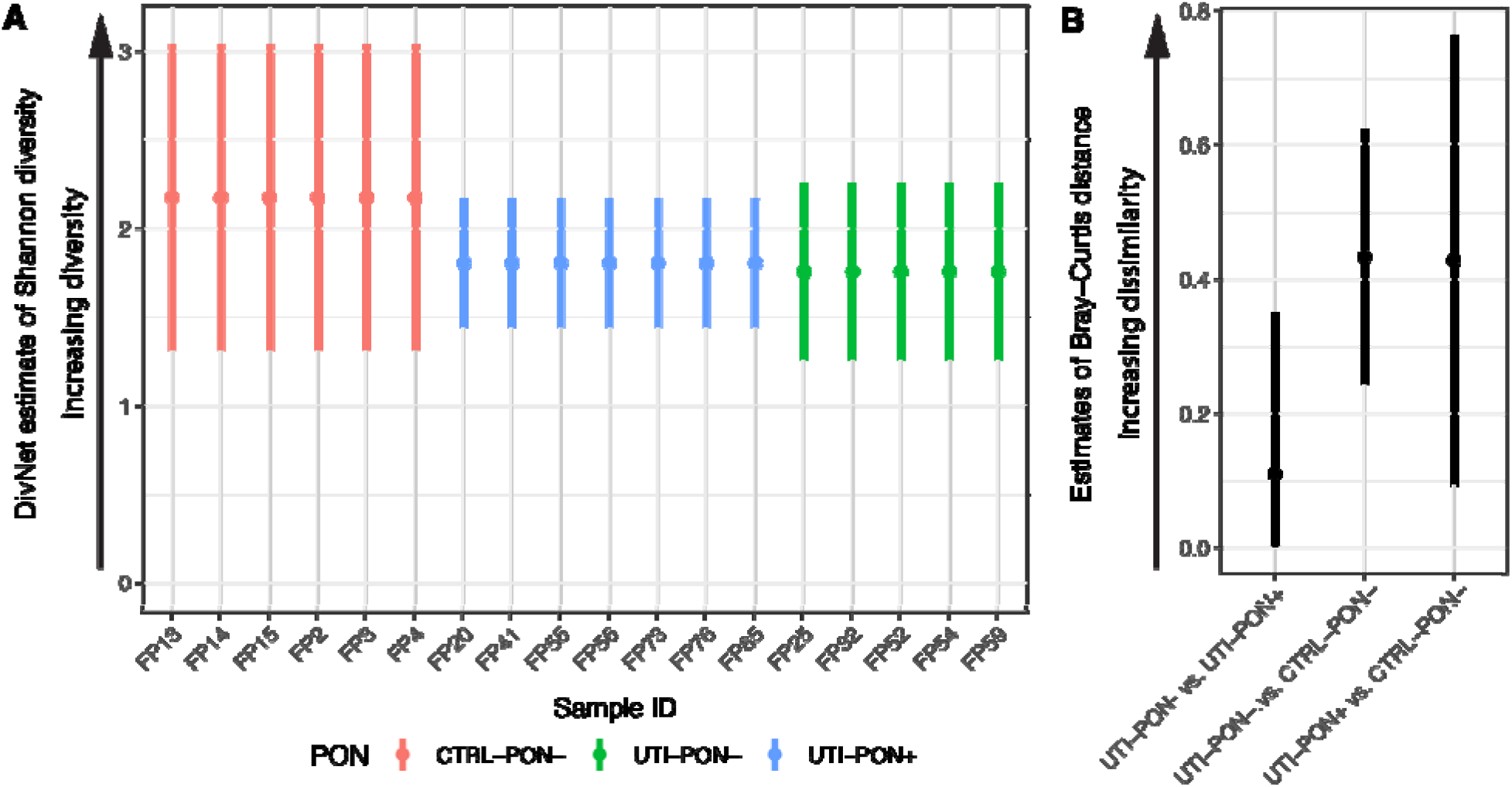
(A) Differences in Shannon diversity estimates (with confidence intervals) were seen between control and UTI subjects, but not between PON+ and PON− UTI subjects (breakaway betta test, p-value < 0.05) (B) Beta diversity estimates (with confidence intervals) highlight that control and UTI samples are more dissimilar to each other than PON+ and PON− UTI samples.

**Figure 3.**
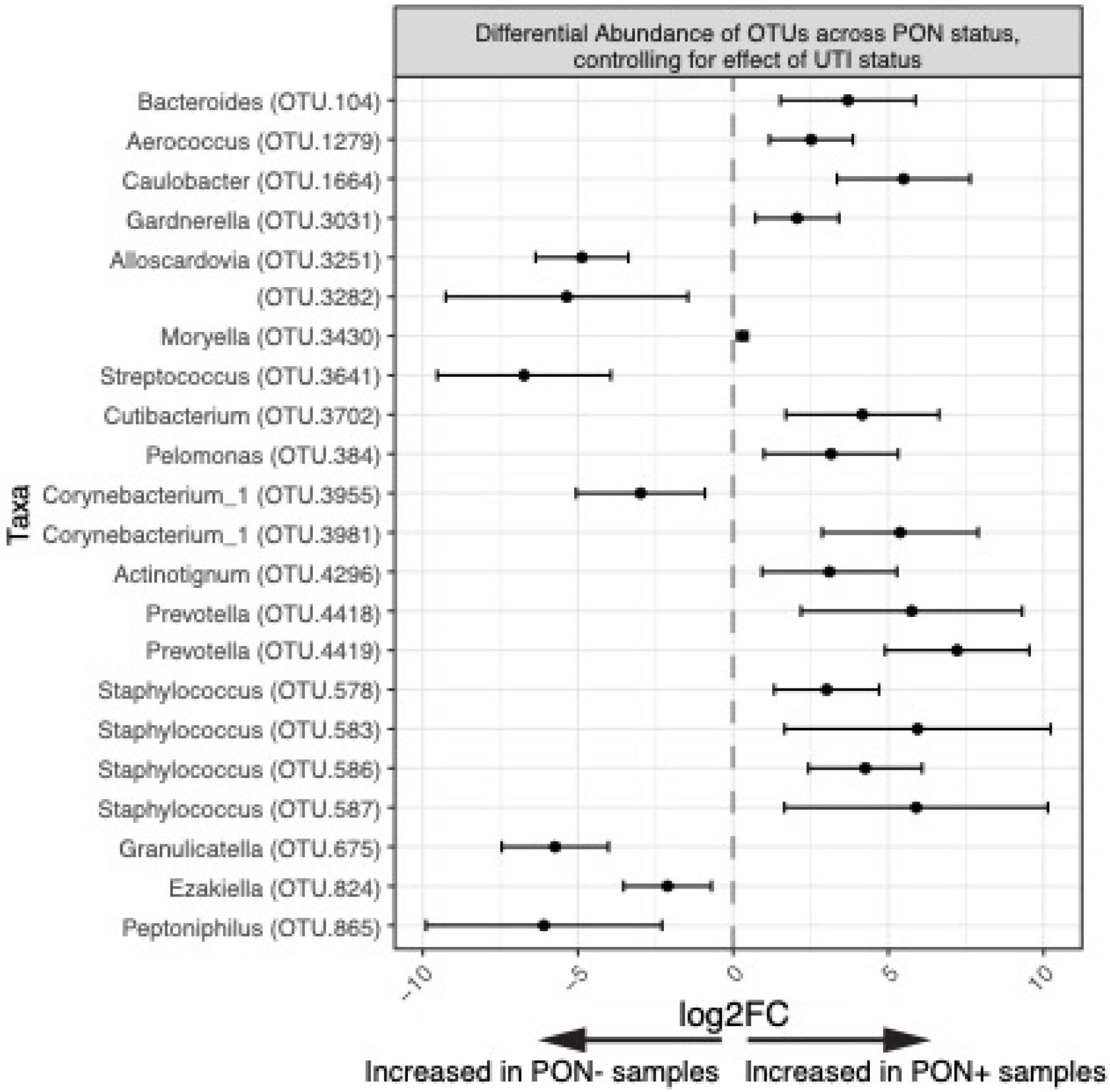
A total of 22 OTUs were differentially abundant across PON status when controlling for the effect of UTI status on abundance. OTUs detected by differentialTest in corncob (FDR adjusted p-values < 0.05). The seven OTUs increased in PON− samples are all small, gram positive host commensals not otherwise found in the environment. Species over-represented in PON+ samples include environmental opportunists such as *Aerococcus* and *Caulobacter*, as well as human-associated genera such as *Bacteroides*, *Gardenerella*, *Cutibacterium*, *Prevotella, and Staphylococcus*

## Discussion

PON has previously been shown to inhibit virulence in certain gram-negative pathogens, and to influence the composition of host microbial consortia. We speculated that PON may have a role in the innate immune response in UTI. Our results indicate a positive association between urine PON and positive culture in patients with UTI. This finding tends to support the idea that PON may be induced by uropathogens, and be associated with infections with uropathogens that grow out in culture. Absence of PON, on the other hand is associated with urine from those without UTI, or, with symptoms of UTI, but with cultures showing “urogenital flora” (Supplemental Table 2) or, “no growth”. The latter two are urine culture results which are not considered to represent significant infection. It is notable that in PON+ samples OTUs that were significantly increased, (compared to PON− samples) were more diverse and contained environmental/opportunistic species (Figure 3), while OTUs differentially found in PON− samples were a more uniform set of commensals. More generally, the PON+ UTI microbiota was different than the PON− UTI microbiota—whether this is a result of the presence of PON or PON expression resulted from pre-existing consortial differences cannot at present be determined. Consortial differences caused by an AHL quorum quenching enzyme related to PON (SsoPox) was recently reported by Schwab, et al. (41). They found that in a complex microbial community the addition of an AHL-degrading enzyme inhibited biofilm formation (even among genera that neither sense nor produce AHLs) and altered the composition of microbial consortia without changing overall community diversity. Combined with the present results, Schwab et al.’s findings suggest the possibility that interfering with AHL signaling can have far-reaching effects on complex microbial communities beyond those limited to specific effects on species that quorum sense with, or have receptors for, AHLs. In connection with this it is notable that only PON− urines grew out *P. aeruginosa* or were found to contain AHLs. This may be a specific effect of PON; there is also evidence that it has more global effects; a practical example is that, according to our findings, in the complex system consisting of host, uropathogen and urinary microbiota, information about PON, heart rate, and white blood count can predict the probability of positive urine cultures.

In our study PON positive subjects had significantly more UTIs caused by *E. coli* alone, rather than multi-species infections, or infections with opportunists such as *P. aeruginosa.* It has previously been reported that the large majority of uncomplicated UTIs in normal hosts are caused by single species (28). Infection associated with impaired immunity is characterized by difficult to eradicate biofilms, polymicrobial infections, and infection with opportunistic organisms that don’t readily infect immune-competent hosts. PON positive subjects featured microbiota with a larger proportion of opportunists, but greater likelihood of single-pathogen urine cultures, suggesting that PON may contribute to immunocompetence. PON− urines grew out more environmental opportunists in culture, compared to PON+ urines, (which grew out none as the primary uropathogen). As noted, the microbiota of PON positive urine in patients with UTI contained a larger proportion of environmental/opportunistic bacteria; whether or not the presence of opportunistic bacteria induces PON (which may then limit their ability to become primary uropathogen) can’t be determined at present. Nonetheless, the role of PON in innate immunity of the airway and skin (9, 20, 60), suggests that urinary PON may also have a protective role. This may occur through degrading the virulence-associated AHLs of some uropathogens, and independently as a mediator of inflammation. Another possible protective effective of PON relates to its putative effects on microbial consortia, and the possibility that host benefits are realized as a result.

We also report here for the first time AHLs in urine from subjects with UTI. C12 AHL levels in urine from subjects with *P. aeruginosa* UTI have not previously been reported, though detection in urine of non-AHL *P. aeruginosa* mediators of QS associated with pulmonary infection has recently been reported (61). In the current study, 3 out of 4 urine samples from which *Pseudomonas* grew out in culture were found to have detectable levels of C12 AHLs. Two of three were below 1 micromolar (Table 5.) Biologically relevant concentrations of AHLs for QS are considered to be 1-5 micromolar (26), and levels of C12 AHL in planktonic cultures necessary to initiate QS-related *las*B expression have previously been reported to be about 1 micromolar (62). This is a concentration of C12 AHLs that is not uncommonly seen in planktonic cultures of *P. aeruginosa.* By contrast, C12 AHL levels associated with *P. aeruginosa* biofilms in flow cells have been found to be hundreds of times higher (63). One possible interpretation of concentrations of C12 considerably below this in three of four samples suggests that QS and virulence expression in *P. aeruginosa* UTI is not a planktonic phenomenon in the urine. QS may be occurring on mucosal surfaces of the bladder/urinary system with bacterial surface colonization where local concentrations of metabolites such as AHLs are likely to be much higher (64), and interactions with mediators of host immunity more intense (65). Urine has recently been reported to independently promote *P. aeruginosa* biofilm formation (66), suggesting that even in the absence of the normal QS-mediated mechanisms for biofilm formation, a biofilm may still be formed in *P. aeruginosa* UTI.

## Conclusion

We report for the first time AHLs in the urine of subjects with *P. aeruginosa* UTIs; the significance of this, and the role that AHLs play in QS among planktonic *P. aeruginosa* remains to be investigated; our finding suggests that QS mechanisms may affect UTI-related microbial consortia, and possibly microbial pathogenesis in UTI. We found that UTI subjects with PON positive urines were much more likely to have uncomplicated *E. coli* UTI. The presence of PON was associated with distinct microbial consortia in which differentially over-represented genera were more diverse, and included opportunistic environmental species. Future work may address the question of whether PON is induced in circumstances in which uropathogens might otherwise establish difficult to eradicate polymicrobial infections more often seen in immunocompromised hosts.

