## Supplemental material for "Paraoxonase and acylated homoserine lactones in urine from patients with urinary tract infections-- relationship to microbial diversity by 16S rRNA gene sequencing"

**Supplemental Table 1.** Association of Positive Culture with PON1 and other patient variables, in patients with UTI.
N and column-% are shown.

| **Culture** | **Culture Neg (n=22)** | **Culture Pos (n=39)** | **p** |
| --- | --- | --- | --- |
| PON1 Positive | 4 (18%) | 18 (46%) | .03 |
| Age | 60 ± 24 | 64 ± 20 | .45 |
| Race  Black  White  Other | 1 (5%)  16 (73%)  5 (23%) | 9 (23%)  23 (59%)  7 (18%) | .17 |
| Gender female | 16 (73%) | 28 (72%) | .94 |
| Catheterized | 1 (5%) | 10 (26%) | .045 |
| WBC | 9.8 ± 3.5 | 12.2 ± 4.6 | .046 |
| Hemoglobin | 12.7 ± 1.7 | 12.3 ± 2.0 | .44 |
| Serum creatinine | 1.1 ± 0.6 | 1.2 ± 1.0 | .44 |
| Highest HR | 86 ± 16 | 97 ± 22 | .064 |
| Highest temp | 98.8 ± 1.4 | 99.0 ± 1.5 | .61 |
| Lowest systolic bp | 119 ± 19 | 116 ± 21 | .58 |
| Lowest diastolic bp | 71 ± 9 | 67 ± 14 | .23 |

**Supplemental Table 2**. Univariable association of presence of PON in urine with *E. coli* alone versus cultures growing out multiple different bacteria (including ‘urogenital flora’)

|  | ***E. coli* alone** (n=21) | **Mult. organisms**.  (n=25) |
| --- | --- | --- |
| PON pos. | 10 (53%) | 6 (24%) |
| PON neg. | 9 (47%) | 19 (76%) |

^A^ Chi-square 5.3; P=0.05

**Supplemental Table 3**. Subjects whose cultures grew out *P. aeruginosa* had urine that contained C12 AHL (except for subject #61 where it was not detectable).

| **Subject** # | **Luminescence** (ALU) | **Micromolar conc**. |
| --- | --- | --- |
| 9 | 167,000 | 1.5 |
| 25 | 22,000 | 0.2 |
| 35 | 8,000 | 0.07 |
| 61 | nd | nd |
